# Chronic distress in male mice impairs motivation compromising both effort and reward processing with altered anterior insular cortex and basolateral amygdala neural activation

**DOI:** 10.1101/2021.05.28.446144

**Authors:** Lidia Cabeza, Bahrie Ramadan, Stephanie Cramoisy, Christophe Houdayer, Emmanuel Haffen, Pierre-Yves Risold, Dominique Fellmann, Yvan Peterschmitt

## Abstract

In humans and mammals, effort-based decision-making for monetary or food rewards paradigms contribute to the study of adaptive goal-directed behaviours acquired through reinforcement learning. Chronic distress modelled by repeated exposure to glucocorticoids in rodents induces suboptimal decision-making under uncertainty by impinging on instrumental acquisition and prompting negative valence behaviours. In order to further disentangle the motivational tenets of adaptive decision-making, this study addressed the consequences of enduring distress on relevant effort and reward processing dimensions. Experimentally, appetitive and consummatory components of motivation were evaluated in adult C57BL/6JRj male mice experiencing chronic distress induced by oral corticosterone (CORT), using multiple complementary discrete behavioural tests. Behavioural data (from Novelty Supressed Feeding, operant effort-based choice, Free Feeding and Sucrose Preference tasks) collectively show that behavioural initiation, effort allocation and hedonic appreciation and valuation are altered in mice exposed to several weeks of oral CORT treatment. Additionally, data analysis from FosB immunohistochemical processing of postmortem brain samples highlight a CORT-dependent dampening of neural activation in the anterior insular cortex (aIC) and basolateral amygdala (BLA), key telencephalic brain regions involved in cue appetitive and consummatory motivational processing. Combined, these results suggest that chronic distress-induced irregular aIC and BLA neural activations with reduced effort production and attenuated reward value processing during reinforcement-based instrumental learning could result in maladaptive decision-making under uncertainty. The current study further illustrates how the stoichiometry of effort and reward processing contributes to dynamically adjust the motivational threshold triggering goal-directed behaviours in versatile environments.

## INTRODUCTION

Motivation is defined as “a reason or reasons for acting or behaving in a particular way” (Husain and Roiser, 2018), and is often understood as a measure of the amount of energy or other resources that an individual is willing to invest towards achieving a valued outcome (Berridge, 2004). Motivation is therefore considered a critical modulator of reward-related behaviour, which may impact high order functioning including decision-making. Individuals’ motivational state therefore modulates their engagement in rewarding behaviours, from initiation (*appetitive behaviour*) to cessation with maintenance through persistent effort (Husain and Roiser, 2018; Salamone et al., 2018).

Apathy, or loss of motivation in goal-directed behaviour has been described as a core symptom across psychiatric disorders subsequent to chronic distress (Pedersen et al., 2009; Hartmann et al., 2015; Husain and Roiser, 2018) including depression (Treadway and Zald, 2011; Treadway et al., 2012). Depressed patients, even if able to experience pleasure *per se*, are usually reluctant to expend effort in exchange for rewarding experiences. In combination with energy-related deficiencies or *anergia*, or even with fatigue, over-estimation of the effort necessary for obtaining rewards and reduced reward anticipation could be at the origin of motivational deficits in this clinical population, potentially leading to a suboptimal integration of cost/benefit information (Treadway and Zald, 2011). Consequently, disruptions in reward processing influence the establishment of adaptive action-outcome associations that lead to the development of interest, desire and anticipation for the latter (Kring and Barch, 2014; Rizvi et al., 2016). Impaired associative learning reported in psychiatric conditions (Horan et al., 2006; Der-Avakian and Markou, 2012; Griffiths et al., 2014) might result from *abulia* (i.e. reduced ability to initiate an action) or *anhedonia* (i.e. reduced ability to feel pleasure or lack of reactivity to positive experience from reward), but they are likely not the only contributors (Pizzagalli, 2014; Husain and Roiser, 2018). Therefore, reward anticipation, which involves momentary motivational arousal in healthy conditions, effort valuation to obtain a reward and motivation in a broad picture are, among others, processes involved in hedonic appreciation that may lead to variability in goal-directed behavioural outputs.

Behavioural adjustment to novel situations is necessary in order to adaptively modify behaviour in dynamic environments, and requires information updating and monitoring necessary for effective cue processing and strategy definition (Seu et al., 2009; Dajani and Uddin, 2015; Uddin, 2021). Consequently, deficits in these processes considerably impact individuals’ ability to learn from the outcomes of their actions and might decrease the quality of their decisions.

Preclinical studies on the effect of chronic distress have evidenced motivational deficits via appetitive and effort-related tasks, and in particular chronic glucocorticoids (GC) administration has been demonstrated to interfere rodents’ instrumental conditioned responding to food reward (Gourley et al., 2008; Olausson et al., 2013; Dieterich et al., 2019, 2020). However, these alterations seem to be highly dependent upon type, intensity and duration of the stressors (Hurtubise and Howland, 2017). Chronic GC administration in mice efficiently induces negative valence behaviours reminiscent to those subsequent to human chronic distress (Dieterich et al., 2019; Cabeza et al., 2021). Yet, results are controversial concerning the impairment of positive valence behaviours (David et al., 2009; Olausson et al., 2013; Dieterich et al., 2019; Cabeza et al., 2021). However, recent results from our laboratory have demonstrated that chronic exposure to GC disrupts male mice decision-making in a valued-based gambling task, mostly by obstructing the exploration-exploitation trade-off.

Several brain regions and networks are suggested to be involved in decisional processing under distress. Particularly, the medial prefrontal cortex (mPFC) and the amygdala have been related to gambling performance in rodents (de Visser et al., 2011), but the convoluted relationship between the phenotype resulting from long-lasting exposure to circulating GC and motivation remains elusive. The present study aims at disentangling the neural underpinnings of the relationship between chronic GC exposure and motivational deficits, in order to obtain insight into the behavioural and neural mechanisms underlying suboptimal decision-making upon distress. For that, the neural activation pattern of multiple discrete brain areas has been explored in relation to appetitive and consummatory motivational behaviours. We forecasted aberrant neural activation patterns in mice after chronic GC administration especially in (i) the mPFC (prelimbic and infralimbic cortices), given its role in instrumental acquisition and appetitive behaviours, for their contribution in reward anticipation and processing, and for their participation in the stress response (Delatour and Gisquet-Verrier, 2000; Akana et al., 2001; Capuzzo and Floresco, 2020); (ii) we also expected the orbitofrontal cortex (OFC) to play a role since it directly participates in effort modulation, and also for its contribution to subjective outcome valuation and contextual salient information integration (Stalnaker et al., 2009; Gourley et al., 2010); (iii) we prognosticated aberrant neural activity within the nucleus accumbens (NAc) – basolateral amygdala (BLA) – ventral tegmental area (VTA) network, which participates in multiple aspects of motivational processing, including reward processing and integration of incentive value dynamics (Wang, 2005b; Roozendaal et al., 2009; Floresco, 2015; Wassum and Izquierdo, 2015; Piantadosi et al., 2017; Bouarab et al., 2019); and (iv) we expected to evidence a significant contribution of the anterior insular cortex (aIC), given its role in interoceptive information and effort-based processing, as well as for its contribution to the representation of anticipatory cues and decision-making (Nieuwenhuys, 2012; Daniel et al., 2017; Gogolla, 2017).

Since the emergence of several neuropsychiatric disorders has been related to the deleterious effects of sustained distress, understanding the underlying mechanisms of the co-occurring motivational deficits may help identifying predictive biomarkers for treatment selection towards precision medicine in biological psychiatry.

## MATERIALS AND METHODS

### Animals

Hundred and twenty-two 6-8 week-old male C57BL/6JRj mice (*EtsJanvier Labs*, Saint-Berthevin, France) were group-housed and maintained under a normal 12-hour light/dark cycle with constant temperature (22±2°C). They had access to standard chow (Kliba Nafag 3430PMS10, *Serlab*, CH-4303 Kaiserau, Germany) *ad libitum* up to one week before the start of the behavioural evaluations, and from then onwards they were under food deprivation to 80-90% of their free-feeding weight (mean ± SEM (g) = 26.33±0.20). Bottles containing water and/or treatment were available at all times.

Experiments were all conducted following the standards of the Ethical Committee in Animal Experimentation from Besançon (CEBEA-58; A-25-056-2). All efforts were made to minimize animal suffering during the experiments according to the Directive from the European Council at 22^nd^ of September 2010 (2010/63/EU).

### Pharmacological treatment

Behavioural assessments started from the sixth week of treatment. Half the individuals received corticosterone (CORT, -4-Pregnene-11β-diol-3,20-dione-21-dione, *Sigma-Aldrich*, France) dissolved in vehicle (VEH, 0.45% hydroxypropyl-β-cyclodextrin -βCD, *Roquette GmbH*, France) in the drinking water (35μg/ml equivalent to 5 mg/kg/day, CORT group, n=61) throughout the entire experiment. The other half of the animals received vehicle in the drinking water (VEH group). Bottles containing CORT or VEH were prepared twice a week (David et al., 2009; Mekiri et al., 2017).

### Behavioural characterization

Behavioural evaluations were conducted during the light phase of the cycle (from 8:00 a.m.). Animals were randomly separated in two groups (A and B) in order to evaluate different motivational components at similar times of treatment. The group A comprised 42 animals (VEH, n=21; CORT, n=21), which were first tested in the Novelty Suppressed Feeding (NSF) task, and then in an operant Progressive Ratio (PR) schedule of reinforcement task. Animals from the group B (VEH, n=40; CORT, n=40) were tested in the Free Feeding Task (FFT) and the Sucrose Preference Test (SPT). Different foods were used as rewards: standard chow was used in the NSF task; 20 mg grain-based pellets (Dustless Precision Pellets® Grain-Based Diet, *PHYMEP s*.*a*.*r*.*L*., Paris, France) were used in the PR task and the FFT; ∼20 mg chocolate pellets (Choco Pops, *Kellogg’s*®) were also presented during the FFT; 2.5% and 0.8% sucrose solutions (D(+)Saccharose, *Carl Roth*, Karlsruhe, Germany) were used for the SPT.

### Fur Coat State (FCS) evaluation

The state of the fur coat frequently degrades in rodents subjected to sustained distress. Its aspect is frequently used as an index of the quality of animals’ self-oriented behaviour and practice of cleaning (Nollet et al., 2013), but scientific evidence has also reported adverse effects of chronic exposure to exogenous GCs on epidermal homeostasis (Pérez, 2011) and hair growth initiation (Stenn et al., 1993).

As an index of the treatment’s efficacy and the quality of self-oriented behaviour, animals’ coat state was evaluated weekly from the beginning of the experiment. For this, seven body regions were rated (head, neck, abdomen, front legs, hind legs and tail) and a score between 0 and 1 was assigned to each one as described by Nollet et al., 2013 (0: good state; 0.5: coat moderately degraded; 1: bad state). A final score between 0 and 7 was calculated adding all scores.

### Novelty Suppressed Feeding (NSF) task

The NSF task evaluates hyponeophagia in rodents (i.e. the inhibition of feeding produced by a novel environment) and has been proved useful to evaluate the emergence of negative valence behaviour in rodents (Samuels and Hen, 2011). During the sixth week of differential treatment, food deprived animals were individually placed in an opaque cylindrical open-field apparatus (diameter: 47 cm; height: 60 cm), in the centre of which two grain-based pellets were located on a white piece of filter-paper (diameter: 10 cm). The latency of the animals to start eating was recorded and used as an index of appetitive behaviour. The test finished immediately after animals started to eat or when a maximum time of 15 min elapsed. Immediately after the test, animals were transferred to an individual home cage where food was available, and the total amount of food consumed during 5 min was measured.

### Operant habituation and training

Operant behaviour was evaluated from the seventh week of differential treatment, using 5 identical two-hole-operant chambers (*Med Associates*, Hertfordshire, UK), individually housed inside sound-attenuating cubicles with ventilating fans and connected to a PC computer with MedPC-5 software via a *Smart Control* interface cabinet. The MedPC programs used in this study were written in Medstate notation code.

Prior to behavioural evaluation, animals followed a habituation period and an operant training. *Habituation*: animals were introduced during 10 min for 3 consecutive days in the operant chambers, where they could explore an empty and illuminated food receptacle in one side, and two unilluminated identical holes in the opposite side. The third day they could find food in the food receptacle. *Training*: following the habituation sessions, mice were trained to nose-poke in one of the holes on a fixed ratio (FR)-1 schedule of reinforcement, where a single nose-poke elicited the delivery of a food pellet. Only the lit hole was designated as “active” and triggered the reward delivery. The allocation of the active and inactive holes was random for each mouse, except for animals displaying spatial preference during the habituation period (twice as frequent visits to a hole respective to the other). For those, the active hole was selected against their preference. Five-second timeout to the FR-1 was given after a nose-poke, for both, right or wrong choices. During the timeout, additional nose-pokes did not result in reward delivery, giving mice the time to consume the food. Each training session lasted an hour, or until a maximum of 50 rewards was delivered. Animals were considered to acquire a food-maintained operant responding (*acquisition criteria*) when they (1) exhibited a discrimination index of ≥3:1 for the active versus inactive responses, (2) obtaining ≥20 rewards per session (3) over three consecutive sessions. When these acquisition criteria were fulfilled, the schedule was increased to FR-5 for which five active nose-pokes triggered the delivery of one food reinforcer. The FR-5 training lasted 3 consecutive days.

### Progressive Ratio schedule of reinforcement (PR)

Effort allocation was evaluated using an operant PR protocol (adapted from Sharma et al., 2012) that started after completion of the operant training. The ratio schedule for the PR testing was calculated as previously described (Richardson and Roberts, 1996) using the following formula: [5*e*^(*R*\*0.2)]^ − 5, where R is the number of rewards already earned plus 1. The ratios to obtain a food reward are: 1, 2, 4, 6, 9, 12, 15, 20, 25, 32, 40, 50, 62, 77, 95 and so on. The last ratio completed by an animal is its breakpoint (BP). A PR session lasted an hour or ended after 10 min of inactivity (absence of visits to the holes or the food receptacle). The PR testing finished when the number of rewards earned in a session deviated ≤10% for 3 consecutive days.

### Free Feeding Task (FFT)

This behavioural paradigm (modified from Vichaya et al., 2014) allows investigating the contributions of positive valence systems to decision-making by focusing on hedonic appreciation and valuation. From the eighth week of differential treatment, animals were free to access and feed from two different sorts of nourishments, including a sweet option appealing to the animal’s natural preference for sweetness. Subjective valuation of the presented options is addressed by the choice itself (i.e. food reward primarily chosen for consumption), while appetitive behaviour is evaluated from the latency until animals are starting to eat. Subsequently, hedonic appreciation is approached through animals’ consummatory behaviour. Cognitive aspects contributing to behavioural initiation such as behavioural flexibility or risk-taking, are inferred from a variation of the paradigm for food neophobia evaluation (i.e. presentation of one unfamiliar sort of food).

Mice were placed in a transparent cage where two recipients with food pellets were exposed, one with 10 grain-based pellets and the other one with 10 chocolate pellets. The nature of the first pellet chosen, as well as the latency of this first choice and the total number of pellets consumed were noted. The test finished when all pieces of the first choice were eaten, or after a maximal test duration of 5 min. Animals which failed to choose within the assigned time were considered as *undecided*. Mice were tested twice in the FFT, one week apart: at the beginning of the neophobia evaluation, animals were familiar with the grain-based pellets but not with the chocolate option, while during the second evaluation (referred as “re-test”) all animals were familiar with both types of rewards.

### Sucrose Preference Test (SPT)

Hedonic appreciation was addressed by studying the preference over water of 2.5% and 0.8% sucrose solutions of animals VEH/CORT treated during 10 weeks. Founded upon animal’s natural preference for sweetness (Berridge, 1996, 2004), this paradigm allows investigating hedonic aspects of motivated behaviours, also referred as ‘liking phase’ (Husain and Roiser, 2018), through consummatory actions.

The SPT lasted four consecutive nights (one night of forced sucrose consumption and 3 nights of free choice) and was performed as previously described (Cabeza et al., 2021). The sucrose preference was calculated as the percentage of sucrose solution consumption relative to the total liquid intake, during the last session. Two different sucrose concentrations were used in order to prevent potential ceiling effects.

### Animals sacrifice and brain sampling

Twelve mice (VEH, n=6; CORT, n=6) randomly chosen from the behavioral group A were sacrificed 24 hours following the attainment of their BP in the PR testing for FosB immunostaining. Animals were deeply anesthetized with an intraperitoneal injection of Dolethal (1 mL/kg, *Vetoquinol*), then transcardially perfused with 0.9% NaCl followed by ice-cold 4% paraformaldehyde (PFA, *Roth®*, Karlsruhe, Germany) fixative in 0.1 M phosphate buffer pH 7.4. Once extracted, brains were post-fixed overnight in the same fixative at 4 °C and cryoprotected by immersion in a 15% sucrose solution (D(+)-Saccharose, *Roth®*, Karlsruhe, Germany) in 0.1 M phosphate buffer 24 hours at 4 °C. Brains were then frozen by immersion in isopentane (2-méthylbutane, *Roth®*, Karlsruhe, Germany) at -74 °C using a Snap-Frost™ system (*Excilone*, France) and stored at -80 °C until processed. Brains were cut in coronal 30 µm-thick serial sections, collected in a cryoprotector solution (1:1:2 glycerol/ethylene glycol/phosphate buffered saline –PBS; *Roth®*, Karlsruhe, Germany) and stored at -40 °C.

### Immunohistochemistry

Several sections were selected in order to study the following brain structures: mPFC (PL and IL areas), OFC and NAc (core –NacC, and shell -NAcS) at [1.94 - 1.70] anterior to Bregma (aB); rostral aIC at [0.86-0.50] and caudal at [0.26 - 0.02] aB; paraventricular nucleus of the hypothalamus (PVN) and dorsal region of the bed nucleus of the stria terminalis (BNST) at [0.26 - 0.02] aB; amygdala (central -CeA and BLA) at [0.94 - 1.34] posterior to Bregma (pB); parasubthalamic nucleus (PSTN) at [2.06 - 2.30] pB; and VTA at [2.92 - 3.16] pB (according to Franklin and Paxinos, 2008). Once mounted on gelatin-coated slides, sections were exposed to an antigen retrieval method to maximize antigenic site exposure for antibody binding. Sections were immersed for 40 min in 10 mM citrate buffer pH 6 (sodium citrate, *Sigma Aldrich*, Germany) previously heated at 96 °C in a water bath. Then, the buffer was left to cool down at room temperature before removing the sections. After several washing, sections were exposed 15 min to a 0.3% hydrogen peroxide solution in order to block endogenous peroxidase activity, avoiding non-specific background staining. Next, sections were incubated with the primary antibody (*Sc-48*, rabbit anti-FosB, *Santa Cruz Biotechnology*, 1:500) during 48 hours at room temperature. Tissues were then incubated with the secondary HRP antibody (*BA-1000*, goat anti-rabbit Ig, *Vector Laboratories*, 1:1500) during 24 hours at room temperature. After washing, the amplification of the signal was addressed by using an avidin horseradish peroxidase complex (ABC Elite kit, *Vector Laboratories*) for 40 min at room temperature. A 3,3′-Diaminobenzidine (DAB) chromogen solution was used to visualize the peroxidase complex. Brain sections were then dehydrated with successive alcohol baths (70°, 95°, 100° and 100°, 3 min), cleared with xylene (*Avantor®*, 3 x 5 min) and finally coverslipped with Canada Balsam (*Roth®*, Karlsruhe, Germany). Microscopic brain images from enzymatic immunohistochemistry were acquired using 4x objectives of an Olympus microscope Bx51 equipped with a camera Olympus DP50. *ImageJ* software was used to count FosB labelled cell nuclei over the regions of interest. Due to technical issues, some sections and related regions could not be included in the final analysis, so that final samples sizes were: PL n=12, IL n=12, OFC n=12, NAc n=12, rostral aIC n=12, caudal aIC n=11, dorsal BNST n=12, PVN n=11, amygdala n=12, PSTN n=10 and VTA n=11.

### Data and statistical analyses

#### Data are presented as means ± SEM

Statistical analyses were conducted using STATISTICA 10 (*Statsoft*, Palo Alto, USA) and figures were designed using GraphPad Prism 9 software (*GraphPad Inc*., San Diego, USA). The samples sizes were identified *a priori* by statistical power analysis (*G*Power* software, Heinrich Heine Universität, Dusseldorf, Germany). Our behavioural sample is predicted to yield highly reproducible outcomes with 1-β>0.8 and α<0.05. Assumptions for parametric analysis were verified prior to each analysis: normality of distribution with Shapiro-Wilk, homogeneity of variance with Leven’s and sphericity with Mauchly’s tests Behavioural and immunohistological scores from the pharmacological conditions were compared by Student t-tests, and when datasets did not meet assumptions for parametric analyses, Mann Whitney U and Wilcoxon tests were used.

Survival curves for NSF scores were compared using the Gehan-Breslow-Wilcoxon (GBW-Χ^2^) method. The assumption of independent and normal distribution of treatment populations within FFT clusters based on differential choice was tested with Chi-squared tests (Χ^2^). Dimensional relationships between behavioural scores and FosB expression in the various brain structures under investigation were analysed using Pearson correlations. For all analyses, the significance level was p<0.05 and analysis with p≤0.1 were described as trends. Effect sizes are reported as partial η^2^ (pη^2^).

## RESULTS

Motivational dimensions potentially impacting decision-making under distress were addressed through various behavioural evaluations. The weekly evaluation of the animals’ FCS served informing about the pharmacological efficacy (pharmacodynamics) and also their self-oriented behaviour (Nollet et al., 2013). Along with the literature, the FCS of CORT-treated mice (n=61) appeared significantly degraded from the third week of treatment compared to the control group (n=61) [main effect of treatment: F_1,1279_=454.8, p<0.00, pη^2^=0.26; week: F_13,1279_=25.0, p<0.00, pη^2^=0.20; treatment x week interaction: F_13,1279_=14.3, p<0.00, pη^2^=0.13; from the third week of treatment: all ps<0.01, post-hoc].

*Group A*. The NSF task was used to study the appetitive dimension of motivation by approaching behavioural initiation to feed. Mice treated with CORT during 6 weeks approached and started eating from the presented food after a significantly longer latency period (n=21; mean latency to eat (s) ± SEM: 537.27±69.74) than VEH-treated animals (n=21; 85.63±11.89) [UMW: Z=-4.8, p=0.0000; GBW, X^2^=28.0, p<0.0001]. Distress mice therefore displayed a typical negative valence behaviour in the NSF task in accord with the literature (David et al., 2009; Dieterich et al., 2019). Of note, the comparison of the food intake after the test, frequently used as appetite control between groups (David et al., 2009), shows an effect of the treatment, with CORT-treated animals eating significantly less (mean food intake (g) ± SEM: 0.09±0.02) than VEH animals (0.04±0.02) [Z=3.9, p<0.0001]. Since the body weight was monitored for all animals throughout the food deprivation procedure, this difference can be attributed to CORT-induced metabolic effects preferentially (Wang, 2005a) rather than to differential hunger pressure. See **Figure 1A,B**.

**Figure 1.**
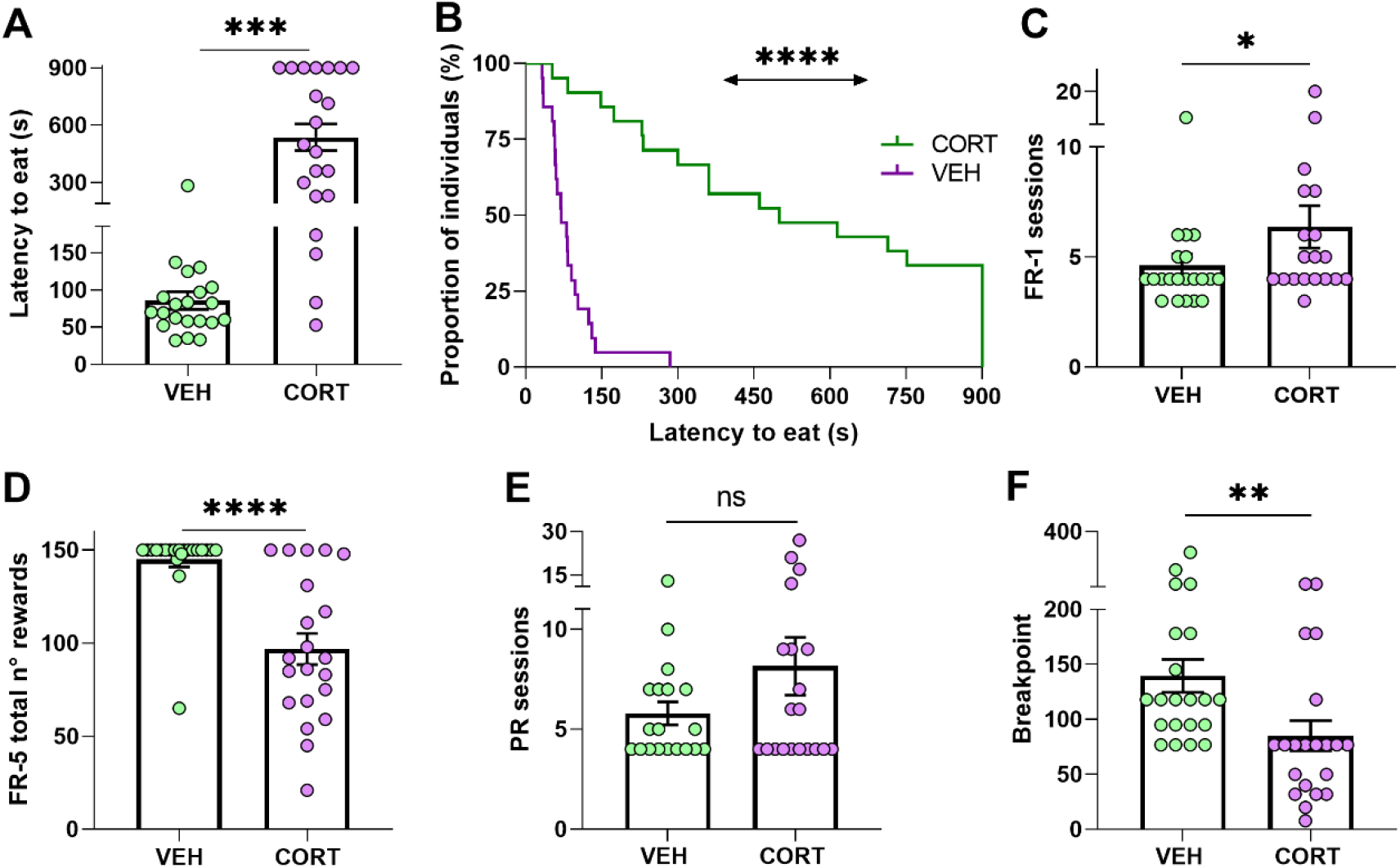
Chronic CORT administration hampers appetitive behaviour in mice. **(A-B)** In the NSF test and under a sustained food restriction protocol, animals chronically treated with CORT took longer to start eating from a food pellet placed in the centre of an open arena (***, p<0.001; ****, p<0.0001), **(C)** Compared to control animals, CORT-treated mice required more sessions to reach the acquisition criteria in the FR-1 reinforcement phase, indicating a detrimental effect of the GC pharmacological treatment in action-outcome association (*, p<0.05). **(D)** During the transitional FR-5 phase, CORT-treated animals obtained significantly less food rewards than control animals (****, p=0.0000). **(E)** All animals, irrespective to their condition, required a similar number of sessions to reach their BP in the PR testing (ns, not significant). **(F)** However, CORT-treated animals significantly allocated less effort to obtain food reward, indicating a deleterious effect of the GC pharmacological treatment on reward and effort valuation and processing (**, p<0.01).

The contingent association of a nose-poke/action – outcome/reward delivery (i.e. instrumental acquisition) was then evaluated from FR-1 data. The comparison revealed that chronic CORT administration resulted in animals requiring more sessions to comply with the acquisition criteria (mean of total number of FR-1 sessions ± SEM, VEH: 4.62±0.51; CORT: 6.37±0.96) [UMW: Z=-2.0, p<0.05] (**Figure 1C**), in agreement with previous studies (see for instance Dieterich et al., 2019).

Preceding the PR testing to study effort allocation, mice behaviour was evaluated in a transitional FR-5 schedule of reinforcement. The quantity of obtained rewards was found to be influenced by the GC pharmacological treatment [main effect of treatment: F_1,40_=26.3, p=0.0000, pη^2^=0.40], with CORT-treated animals (mean number of rewards ± SEM: 96.86±8.45) obtaining a smaller amount of food than controls animals (144.95±4.06) (**Figure 1D**), but without significant effect of the session [session: F_2,80_=1.73, p=0.18, pη^2^=0.04]. CORT- and VEH-treated animals also differed in the percentage of correct responses across the FR-5 reinforcement phase [treatment: F_1,40_=9.7, p<0.01, pη^2^=0.19], though both groups responded almost always correctly (mean percentage of correct responses ± SEM, VEH: 93.84±0.98; CORT: 89.67±0.92). However, these differences need of careful interpretation and are not to be directly attributed to the animals’ motivational state, since intrasession satiation might had influence animals’ behaviour.

The explicit study of motivation to expend effort for a food reinforcer during the PR testing revealed a significant deleterious effect of the pharmacological treatment, again in line with previous studies (Olausson et al., 2013; Dieterich et al., 2019). Animals were tested daily on the PR schedule of reinforcement until they stabilized their performances, i.e. they reached their BP, and no difference between pharmacological conditions was found for the number of required sessions (mean number of sessions ± SEM, VEH: 5.79±0.57; CORT: 8.15±1.45) [UMW, Z=-0.6, p=0.57] (**Figure 1E**). Congruently with the scientific literature, these results showed that mice chronically treated with CORT had a lower BP in the PR task (mean BP ± SEM, CORT: 85.33±13.62) compared to animals from the control group (VEH: 139.57±14.96) (**Figure 1F**). In other words, chronic CORT administration lessened the animals’ motivation to allocate effort for food rewards, in agreement with previous findings on altered reward and effort valuation (see for instance Gourley et al., 2008).

*Group B*. The FFT paradigm used to assess reward appreciation allowed to simultaneously address the influence of food neophobia in mice (VEH, n=40; CORT, n=40). The results show that the majority of individuals chose preferably the familiar grain-based option (57.5%), while only 12.5% of them chose chocolate pellets. The remaining 30.0% of the animals, the so-called *undecided*, failed to choose between the presented rewards within the 5 testing minutes. Interestingly, 72.5% of VEH and 42.5% of CORT-treated animals chose the grain-based reward. Individuals that chose the unfamiliar, chocolate reward represented 22.5% of the control animals as opposed to only 2.5% of the CORT-treated mice. Noteworthy, 55.0% of CORT-treated animals did not select an option, while only 5.0% of VEH-treated animals failed to choose between options before the end of the task. The distribution of CORT- and VEH-treated mice between the three possible classes was compared, highlighting a significant difference [Χ^2^, p=0.0000] which is mainly accounted for by the chocolate reward and undecided choices (see **Figure 2A**).

**Figure 2.**
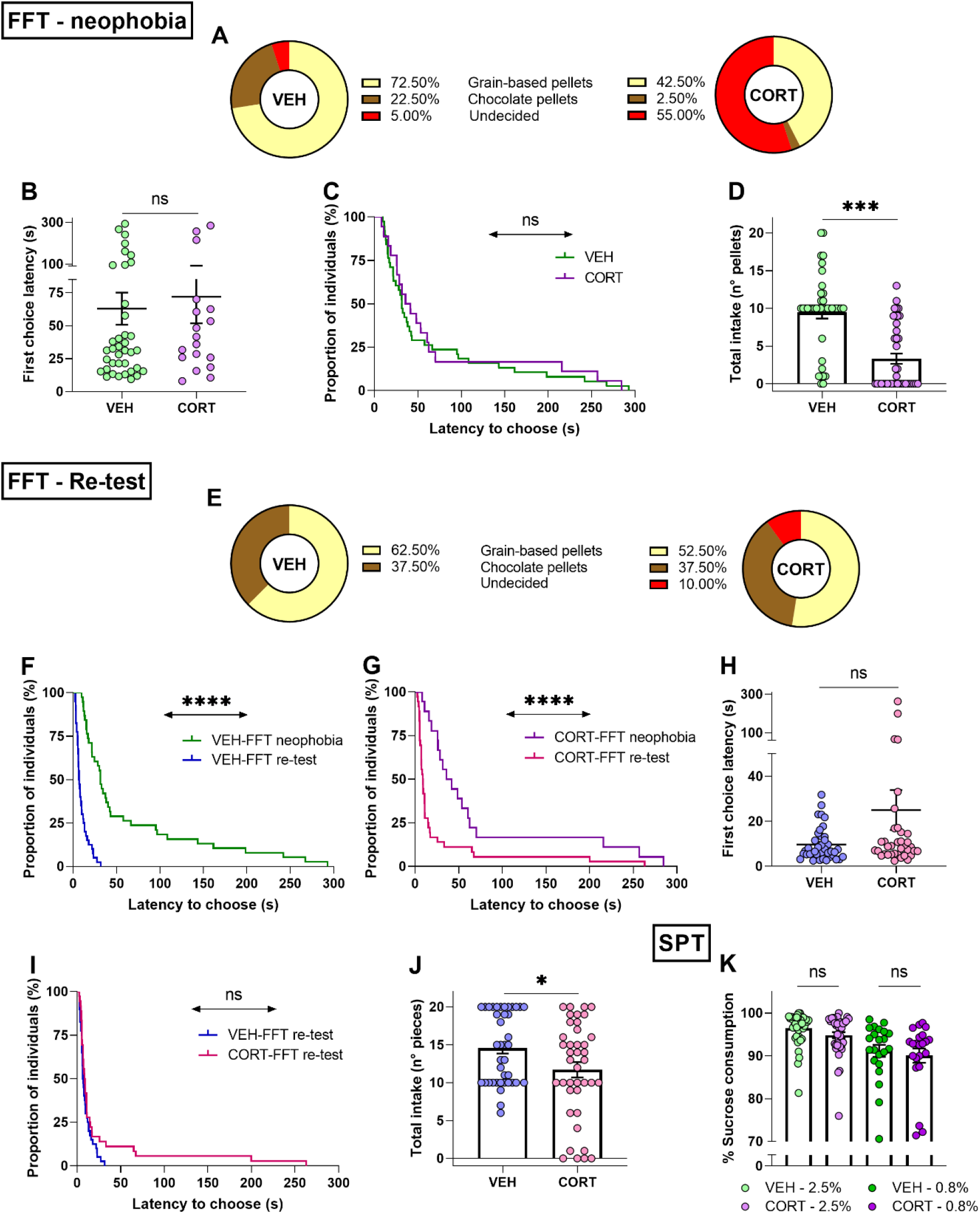
Chronic CORT administration impacts behavioural initiation and consummatory behaviour in mice. **(A) Food neophobia is emphasized after CORT exposure in the FFT.** Different frequency distribution of VEH and CORT-treated individuals in their choice between familiar (grain-based pellets, VEH: n=29; CORT: n=17) and unfamiliar food rewards (chocolate pellets, VEH: n=9; CORT: n=1). Chronic CORT administration affects appetitive behaviour through increment of the proportion of animals failing to choose between rewards within the test duration (undecided, VEH: n=2; CORT, n=22). **(B)** During the neophobia approach of the FFT, VEH and CORT-treated animals needed the same amount of time to make a choice and to start eating. **(C)** Appetitive behaviour of animals that eat (independently of their choice) was not affected by the CORT treatment, as shown by the negligible difference in proportions of individuals starting to eat. **(D)** CORT-treated mice ate significantly less than control animals during the FFT. **(E) Chronic CORT exposure does not affect reward valuation, but impinges on appetitive behaviour in the FFT re-testing**. Different frequency distribution of VEH and CORT-treated individuals in their choice between grain-based pellets (VEH: n=25; CORT: n=21) and chocolate pellets (VEH: n=15; CORT: n=15). Chronic CORT administration affects appetitive behaviour, as evidenced through an increment of the undecided class (undecided, VEH: n=0; CORT, n=4). **(F)** Animals from the control group started eating significantly earlier in the FFT-re-testing **(G)** as well as CORT-treated animals, indicating neophobia overcoming. **(H)** In the FFT re-testing, VEH and CORT-treated animals did not differ in their latency to choose between the two options and start eating (data from undecided individuals excluded). **(I)** VEH and CORT-treated individuals displayed similar appetitive behaviour (independently of their choice), as shown by the lack of difference in the proportion of individuals starting to eat in the FFT re-testing. **(J)** Effect of the CORT treatment on mice consummatory behaviour, with treated animals eating significantly less than control animals. **(K) Chronic CORT administration does not significantly influence hedonic appreciation through consummatory behaviour in the SPT**. Mice chronically treated with CORT showed a significant preference for the sweet solution compared to water [2.5%, n=40, % of sucrose vs 50% solution: t_39_=61.6, p<0.00], as control animals [2.5%, VEH, n=40: t_39_=79.3, p<0.00], and irrespective of the sucrose concentration [0.8%, CORT, n=22: Z=4.1 p<0.0001; VEH, n=22: Z=4.1, p<0.0001] (ns, not significant; *, p<0.05; ***, p<0.001; ****, p<0.0001).

Along the same line, the latency of the first choice was compared between the different conditions, showing that VEH-treated (n=38) and CORT-treated (n=18) animals, if they chose, required a similar amount of time to select their first pellet (latency of the first choice (s) ± SEM, VEH: 62.90±12.17; CORT: 71.93±20.22) [t_54_=-0.4, p=0.69; GBW, X^2^=0.5, p=0.50] (**Figure 2B,C**). Additionally, the total number of eaten pellets as compared between conditions, evidencing a significantly reduced intake in the CORT condition (reward intake ± SEM, VEH: 9.48±0.81; CORT: 3.33±0.68) [t_78_=5.8, p=0.0000] (**Figure 2D**). Combined, these results are consistent with those obtained for the NSF (longer latencies to start eating) and PR (reduced effort allocation) tasks, showing altered appetitive and consummatory behaviour after chronic exposure to CORT. They therefore point to disrupted reward appreciation with CORT-induced exacerbation of the natural food neophobia of rodents.

The same mice (VEH, n=40; CORT, n=40) were re-tested in the FFT paradigm a week after the first behavioural assessment. Confronted to a choice between familiar rewards, the majority of individuals continued to choose in the primarily the grain-based option (57.5%). However, the choice of chocolate pellets increased to 37.5%. Noteworthy, *undecided* represented only 5.0% of the animals. Individuals opting for the non-chocolate reward represented 62.5% of VEH and 52.5% of CORT-treated animals. The chocolate reward was chosen by 37.5% of VEH and CORT-treated mice. Notably, the *undecided* group was only comprised by CORT-treated animals. As previously, the distribution of individuals of the CORT and VEH conditions into the three possible classes was compared. In this re-testing phase, CORT-treated animals chose the chocolate reward as frequently as control animals, but the distribution of the *undecided* group was significantly different compared to the results of the neophobic testing [Χ^2^, p<0.05] (**Figure 2E**).

Regarding the latency to choose, individuals from both conditions required less time to select their first reward compared to the first testing session [FFT-neophobia vs FFT-re-test, VEH: t_37_=4.5, p<0.0001; CORT: t_17_=3.2, p<0.01; GBW, VEH: X^2^=47.8, p<0.0001; CORT: X^2^=16.3, p<0.0001] (**Figure 2F,G**). When compared, CORT-treated animals (n=36) required longer than VEH-treated animals (n=40) to select their first reward, although the difference is not significant (latency of the first choice (s) ± SEM, VEH: 9.64±1.14; CORT: 25.02±8.89) [t_74_=-1.8, p=0.07; GBW, X^2^=1.3, p=0.25] (**Figure 2H,I**). Furthermore, the total food intake over the task continued being low in CORT-treated animals (reward intake ± SEM, VEH: 14.58±0.74; CORT: 11.70±1.01) [t_78_=2.3, p<0.05] (**Figure 2J**).

Hedonic appreciation in mice was then evaluated through their consummatory behaviour in the SPT. The corresponding results demonstrate that all individuals expressed a strong preference for the sucrose solutions compared to water, independent of the sucrose concentration [consumption of sucrose solution vs 50%: all ps<0.0001]. No differences were found between CORT-treated mice and individuals from the control group concerning their consummatory behaviour, independently of the concentration of the sucrose solution [2.5% sucrose solution: t_78_=1.8, p=0.08; 0.8% sucrose solution, UMW: Z=0.5, p=0.60] (see **Figure 2K**). Hence, the SPT did not evidence altered reward processing or hedonic underappreciation in mice after chronic CORT exposure.

Combined, these results do not evidence a significant impairment of reward valuation in mice following chronic CORT administration when food rewards are familiar, but they do indicate an obstruction of behavioural initiation reminiscent to human abulia. Moreover, the reduction of reward intake in CORT-treated animals during the FFT shows an impact of the GC pharmacological treatment on consummatory behaviour, although not confirmed by the SPT evaluation. In order to better understand this difference, the total consumption of sucrose solution (2.5%) and the total intake during the FFT re-test phase were directly confronted. A significant correlation between SPT and FFT scores was obtained [r=0.4274, p<0.0001], indicating that chronic CORT administration might affect consummatory behaviour in mice, and suggesting that SPT assessment at 2.5% might not be adapted to reveal this effect.

Behavioural results were then related to neural patterns of activation in the brain regions of interest, aiming at identifying neural correlates for the CORT-induced behavioural modifications. The levels of chronic neural activation addressed by FosB expression after the PR testing are presented in **Figure 3A**. The density of FosB labelled cells was significantly decreased in the caudal aIC and the BLA of CORT-treated animals (mean of FosB+ cells per mm^2^ ± SEM, caudal aIC: 265.31±6.46; BLA: 603.71±56.07) compared to control individuals (caudal aIC: 369.14±38.85; BLA: 765.78±49.34) [UMW, caudal aIC: Z=2.7, p<0.01; BLA: Z=2.1, p<0.05]. A negative trend of reduced FosB labelled cells density was also evidenced in the dorsal BNST (VEH: 337.41±29.97; CORT: 227.11±29.81) [Z=1.9, p=0.05] and the VTA (VEH: 342.43±26.04; CORT: 249.05±41.85) [Z=1.6, p=0.10] of CORT-treated animals. Otherwise, no differences between conditions were found in the other brain regions here studied [Z<1.6, all ps>0.20]. Nevertheless, the vast SEM values obtained for the CeA of the CORT condition suggest that this dataset should be replicated or focus on anatomical subdivisions (Barbier et al., 2020) in order to circumvent inaccurate interpretation. Therefore, these results indicate an impact of the GC pharmacological treatment on effort and reward valuation and processing during PR testing, by inducing aberrant functional neural activation and plasticity on aIC and BLA respectively.

**Figure 3.**
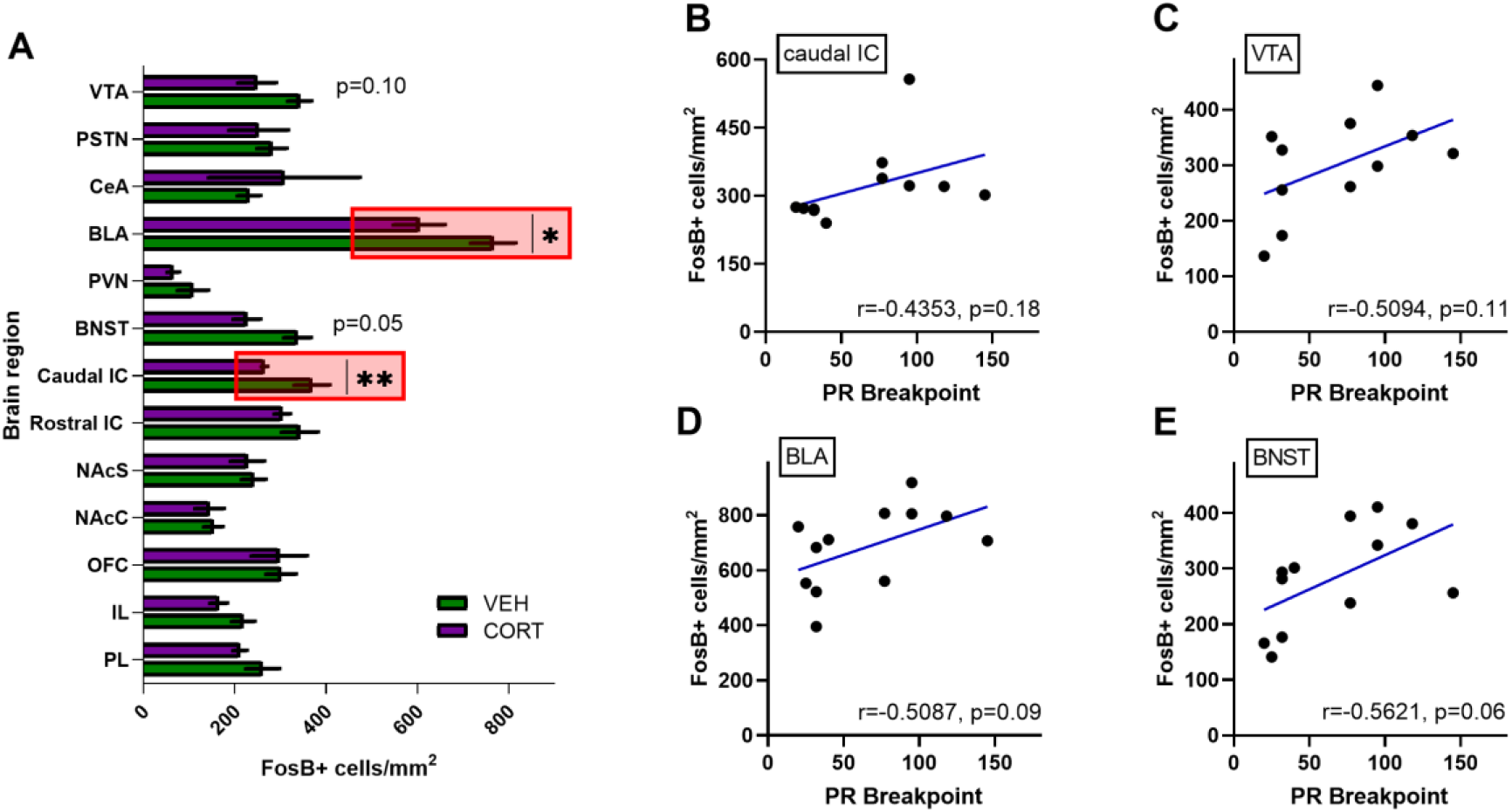
Relative quantification of FosB labelled cells densities in various discrete brain regions of interest following effort-allocation testing in a PR schedule of reinforcement task. **(A)** FosB quantification is expressed as mean of number of immunoreactive labelled cells per surface (mm^2^). Similar activation patterns were found in control and CORT-treated animals in the following brain regions: the PL (VEH: 261.31±36.82; CORT: 211.77±15.14) and IL cortices (VEH: 218.61±25.19; CORT: 165.12±18.75); the OFC (VEH: 301.67±32.81; CORT: 297.65±61.01); the NAcC (VEH: 153.46±21.15; CORT: 145.10±31.73) and NAcS (VEH: 241.737±26.34; CORT: 228.17±36.96); the rostral aIC (VEH: 342.54±39.72; CORT: 304.36±17.50); the PVN (VEH: 108.68±33.85; CORT: 66.02±13.20); the CeA (VEH: 231.37±25.40; CORT: 308.48±165.63); and the PSTN (VEH: 281.70±32.64; CORT: 252.22±64.90). However, a significant effect of the GC pharmacological treatment was evidenced in the caudal aIC (**, p<0.01) and the BLA (*, p<0.05), with reduced activation in CORT-treated animals respect to control individuals. Simultaneously, two negative trends of activation were found in CORT animals in the BNST (p=0.05) and the VTA (p=0.10). Among overall data (VEH + CORT individuals), no significant correlation was found between breakpoint (BP) scores in the PR schedule of reinforcement task and FosB labelled cells densities, irrespective of the brain region considered, **(B)** including the caudal aIC and **(C)** VTA. However, two trends between the behavioural and neurological scores were observed **(C)** in the BLA (p=0.09) and **(E)** in the BNST (p=0.06), pointing to an intricate relationship between neural activation patterns and effort allocation valuation, as major motivational component, in control and stressed animals.

Since a detrimental effect of the GC pharmacological treatment was evidenced on instrumental acquisition in the FR-1 schedule reinforcement (i.e. the number of sessions animals needed to associate correctly responding with reward presentation), FosB labelled cells densities obtained for the different brain regions of interest were confronted with FR-1 behavioural scores aiming to substantiate this finding. However, no significant direct correlations were found between behavioural and neurobiological scores, irrespective of the brain region investigated [r, all ps>0.10].

Similarly, BP scores were systematically confronted with FosB labelled cells densities aiming at directly appraising the neuronal bases of differential animal motivation in the PR paradigm. Despite the fact that no significant correlations were found, including analyses of the caudal aIC [r=0.4353, p=0.18] (**Figure 3B**) and the VTA [r=0.5094, p=0.11] (**Figure 3C**), two positive trends were evidenced for BP scores and FosB labelled cells densities in the BLA [r=0.5087, p=0.09] (**Figure 3C**) and the BNST [r=0.5621, p=0.06] (**Figure 3D**). These results therefore suggest a more intricate relationship between neural activation patterns and effort allocation in the PR task specifically, and point towards the influence of the BLA as a modulator of mice effort-based motivation in physiological conditions and upon chronic distress.

## DISCUSSION

The results of this experimental investigation show that chronic GC exposure has complex detrimental influences on motivational components subserving goal-directed behaviours and specifically, on appetitive and consummatory aspects involved in effort and reward valuation. These motivational deficits likely ground on alterations of interoceptive and sensory cue-related information processing, from neural network computations in somatosensory and associative cortices with neuromodulatory influences of the so-called “brain reward system”.

The present investigation studied various complementary motivational components through food rewarding tasks allowing to address hedonic valuation and appreciation of the reinforcers. Altered appetitive behaviour was observed in distressed animals, as shown by the increased latency to feed in the NSF task, the decreased effort allocation in the PR task and the increased frequency of *undecided* individuals as assessed in the FFT. These results therefore demonstrate an effect of high circulating GC plasma levels on animals’ ability to initiate behavioural actions, in a similar manner to human abulia (Husain and Roiser, 2018). Besides, this could also reflect psychomotor deficits reminiscent to those observed in neuropsychiatric conditions in accord with our previous preclinical (Cabeza et al., 2021) and clinical data (Bennabi et al., 2013). Thus, this result adds value to data suggesting that chronic CORT exposure yields mice especially vulnerable to action-outcome encoding and further consolidation in food reward-based tasks, in accord with our previous results on value-based decision-making under uncertainty (Cabeza et al., 2021).

The study of positive valence-based motivated behaviours in rodents is valuable for translational research in reinforcement learning, but unfortunately literature is incongruent with regards to findings concerning the CORT model. As chronic CORT administration induces a consistent negative valence phenotype (David et al., 2009a; Dieterich et al., 2019), not all aspects of hedonic appreciation, including reward valuation, are reported altered (Cabeza et al., 2021). Grounded on previous studies showing different consummatory scores in healthy mice displaying different gambling strategies (Pittaras et al., 2016), this study sought for new scientific evidence on distress-induced underappreciation of reward as a modulator of decisional strategies. Chronic GC administration did not differentially impact animals’ reward choice in the FFT, suggesting similar hedonic valuation, but the overall food intake was lower in CORT-treated animals. Besides, SPT scores did not differ across pharmacological conditions. Thus, chronic GC exposure may affect hedonic reward appreciation through restraining initiation of actions, rather than through obstruction of reward valuation. This apparent discrepancy might result from protocol variation, and even though various sucrose concentrations have been used in this study experiments, ceiling effects on palatability cannot be excluded. Further investigations are therefore necessary to better disentangle the contribution of hedonic impact on adaptive decisional capability.

Interestingly, the FFT paradigm, which allows evaluating hedonic appreciation and particularly reward valuation, successfully suited the study of additional behavioural aspects potentially influencing goal-directed behaviours. We demonstrated that chronic GC exposure exacerbates food neophobia in mice and most importantly, it prompts indecision. Animals treated with CORT more frequently vacillated or had difficulty in initiating their choice between food rewards when one option was unfamiliar to them.

The operant-based experimental design used in the present investigation enabled revealing a delayed formation of action-outcome associations in individuals chronically exposed to high circulating levels of GC, in accord with an altered exploration-exploitation trade-off demonstrated in a valued-based decision-making task (Cabeza et al 2021). The CORT-induced dysfunctional amotivated state in animals, as shown by the decrease in effort allocation for reward and which resembles to human apathy, further substantiates these findings.

Through impacting cue processing and valuation, chronic GC exposure might obstruct action-outcome encoding and subsequently alter behavioural adaptability in dynamic environments, yielding to poor operational learning. It is therefore plausible that CORT-induced altered reward/effort valuation, processing and integration will impact reward responsiveness through reinforcement learning in valued-based decisional tasks. Chronic GC exposure would therefore reduce the learning capabilities of mice, potentially hampering the integration of environmental contingencies necessary to long-term beneficial performance.

The discrete “localizationist” approach here used to study the neurobiological underpinnings of CORT-induced motivational deficits revealed a pharmacological impact on neural activation of selected discrete brain regions involved in effort and reward processing (Gogolla, 2017; Beyeler et al., 2018). Anterior IC and BLA neural activations were decreased subsequent to long-lasting exposure to GC. This study used different coordinates within the rodent aIC, in order to address reward-related processing and the representation of anticipatory cues (Kusumoto-Yoshida et al., 2015; Gehrlach et al., 2020) as essential processes of decision-making. Indeed, the IC participates in decision-making under uncertainty (Clark, 2010), and the aIC in particular, is involved in contingency acquisition on rodent gambling tasks by modulating reward preference over punishment (Pushparaj et al., 2015). The IC is also relevant for its contribution in initiating feeding behaviour as a part of a basal ganglia-like network (Barbier et al., 2020), and depending on cognitive and emotional influences. Hence, chronic GC exposure could impact animals’ motivational state by interfering with aIC-related neural network activity, and disrupt cue-processing associated with food delivery in the PR paradigm. The CORT-induced aberrant aIC neuronal activity here identified might reflect an altered anticipatory functioning that curtails reward expectation processing and consequent conditioned responding. However, the lack of significant correlation between BP scores and discrete caudal aIC FosB labelled cells densities pleads for more complex than direct causality relationships.

There is also strong evidence for the prominent role of the BLA in appetitive learning and conditioning by encoding motivational information related to instrumental association, as well as for its contribution during effort-based rewarding tasks (Wassum and Izquierdo, 2015; Beyeler et al., 2018; Ferland et al., 2018). In this regard, chronic GC exposure might impact BLA neural activity, obstructing animals’ motivation to effort allocation for food reward likely by disrupting reward value encoding. Moreover, the positive trend found in the correlation between BP scores and BLA FosB labelled cells densities aligns with this hypothesis and could also explain the trend found for the BNST. This considers the involvement of the unidirectional excitatory BLA-to-BNST projections in behavioural output signalling. The BNST, in addition to its contribution on behavioural signalling, is highly responsive to stress and has been associated to distress-related behavioural dysfunction via *crf* signalling (Gray et al., 1993; Ulrich-Lai and Herman, 2009; Hu et al., 2020). However, further investigation is necessary in order to characterize the neural cell types involved in the activity changes identified. Further clustering analyses based on the neurobiological data will also be useful for identifying neurobiological biomarkers predictive of particular behaviours. Combined, these results further substantiate that CORT-treated animals show a compromised motivational state rooted on aberrant aIC and BLA-related neural networks activations involved in effort and reward processing, which could hamper decision-making processing.

Unexpectedly, the neural activation pattern in the NAc was not significantly impacted by sustained exposure to GC, even if a recent study suggests its causal role in instrumental reward-seeking in the CORT model (Dieterich et al., 2021). Similarly, an stronger effect of the GC pharmacological treatment was expected on the VTA neural activation, since its contribution is considered crucial for the modulation of reward motivated behaviours via associative learning (Bouarab et al., 2019). Ongoing analysis aiming at addressing these points of conflict should help clarifying the contribution of the discrete regions of interest and the networks they are part of, to behavioural adaptability in physiological and distress-related conditions.

Modelling distress conditions in animals results, for instance, in disruption of fronto-striatal cognition (Uribe-Mariño et al., 2016; Hupalo et al., 2019) and compromises positive valence systems (Birnie et al., 2020) through alteration of *crf* signalling. In a previous study we have demonstrated that chronic CORT exposure induces suboptimal decision-making in distressed male mice (Cabeza et al 2021), and we have suggested that this alteration relates to high mPFC CRF levels. Here we demonstrated that chronic CORT also prompts reward and effort processing, and learning deficits in male mice. The results of this investigation are therefore in accord with clinical data on motivational deficits (Darcet et al., 2014; Salamone et al., 2018) and suggest that appetitive aspects of motivated behaviours are key players in the modulation of decision-making, particularly in dynamic environments. However, future studies should examine the chronic CORT-induced effects on motivation and decision-making in female mice, since the GC pharmacological treatment differentially induces the emergence of negative valence behaviours across sexes (Mekiri et al 2017; Yohn et al 2019).

In sum, the presented results provide novel insight into the mechanisms of motivational-based learning impairments induced by chronic exposure to GC, and support the validity of the chronic CORT model for translational investigation on effort and reward processing across species and multiple discrete clinical conditions associated to chronic distress. The results are therefore relevant for understanding the pathophysiological mechanisms underlying amotivated-based behaviours in a transdiagnostic perspective.

## Acknowledgements

The authors thank the Animal Facilities of Besançon for technical support and Prof David Belin (University of Cambridge) for valuable scientific contribution.

## Author contribution

LC, YP and DF conceived the experimental design. LC, BR, SC and CH contributed to the data acquisition. LC and YP analysed the data. LC and YP wrote the manuscript. All authors critically revised the work and approved the version to be published. LC, YP and DF agree to be accountable for all aspects of the work in ensuring that questions related to the accuracy or integrity of any part of the work are appropriately investigated and resolved.

